# Bayesian parameter estimation for phosphate dynamics during hemodialysis

**DOI:** 10.1101/2022.06.16.496370

**Authors:** Katrine O. Bangsgaard, Morten Andersen, James G. Heaf, Johnny T. Ottesen

## Abstract

Hyperphosphatemia in patients with renal failure is associated with increased vascular calcification and mortality. Hemodialysis is a conventional treatment for patients with hyperphosphatemia. Phosphate kinetics during hemodialysis may be described by a diffusion process and modeled by ordinary differential equations. We propose a Bayesian model approach for estimating patient-specific parameters for phosphate kinetics during hemodialysis. The Bayesian approach allows us to both analyze the full parameter space using uncertainty quantification and to compare two types of hemodialysis treatments, the conventional single-pass and the novel multiple-pass treatment. We validate and test our models on synthetic and real data. The results show limited identifiability of the model parameters when only single-pass data are available, and that the Bayesian model greatly reduces the relative standard deviation compared to existing estimates. Moreover, the analysis of the Bayesian models reveal improved estimates with reduced uncertainty when considering consecutive sessions and multiple-pass treatment compared to single-pass treatment.

## 1 Introduction

Phosphate enables the body to perform vital processes such as construction of nucleic acids, energy transport and bone tissue formation [3]. The level of phosphate is tightly controlled, and excess phosphate is excreted by the kidneys [20]. However, for patients with renal failure, the control of phosphate homeostasis is impaired. An abnormal level of phosphate is associated with increased vascular calcification and mortality [5, 16].

About half of all dialysis patients suffer from hyperphosphataemia, and strategies to control phosphate levels include phosphate binders, low-phosphate diet and removal of phosphate by hemodialysis [11]. Hemodialysis (HD) is a conventional treatment for renal failure where a patient is coupled to a dialysis machine for four to eight hours. The blood plasma and dialysate fluid are passed through a filter that causes a diffusion process that removes toxic substances, e.g., phosphate, from the blood to the dialysate. The phosphate kinetics in HD is of particular interest because it differs the other removed toxins, e.g., urea, by the fact that hypophoshataemia is fatal for the patient [25]. Thus, the phosphate concentration should not be exhausted, but kept within the critical values.

### 1.1 Previous studies

The control of the phosphate concentration is a considerable clinical problem and has been studied extensively [1, 2, 5, 6, 7, 8, 12, 17, 18, 24]. The conventional hemodialysis treatment is the single-pass (SP) treatment. Agar et al. [1] and Debowska et al. [5] both study the SP treatment by considering a simple two-compartment ordinary differential equation (ODE) model for phosphate removal during HD. They present their results as an average of the measured patients to obtain confidence intervals for their parameters, however, these are not patient specific. Poleszczuk et al. [20] extend the model proposed by Debowska et al. [5] to include a time delay. The time delay is introduced to improve the fit at the later stage of the HD where a minor rebound is observed in some clinical experiments. Andersen et al. [2] analyze the same model analytically and estimate parameters using an optimization-driven approach. Here the parameters are estimated for each patient, but the uncertainty of the parameter estimates is not addressed. Laursen et al. [17] propose a two- and three-compartment model for phosphate clearance during single pass and find that the three-compartment model produces the most satisfying fit but does not address the uncertainty associated with the parameter estimates. Spalding et al. [24] propose a complicated four-compartment model where the fourth pool is a control pool for avoiding dangerously low phosphate concentrations. They argue that a simple two-compartment model cannot fit the relapse phase sufficiently, however, both Andersen et al. [2] and Debowska et al. [5] demonstrate that the simple two-compartment model can produce adequate fits for the relapse phase as well.

A novel HD treatment called multiple pass (MP) [7, 8, 12] provides an alternative to the conventional SP. This novel treatment reduces the amount of dialysis fluid needed for a single session of HD. Andersen et al. [2] and Heaf et al. [12] analyse and compare the MP treatment and SP treatment.

However, none of the above-listed models address patient specific uncertainties associated with the parameter estimates. Moreover, the reported uncertainty of the parameter estimates for the average of the measured patients is very large, e.g., Debowska et al. [5] report a phosphate clearance with a relative standard deviation of 79% and Agar et al. [1] report a relative standard deviation of 47%, indicating that parameters of the two-compartment model are poorly identified. Common for all models is that they assume that the phosphate concentration in the inner-source compartment is known exactly through measurements at time zero. However, measurements are noisy and can potentially bias the results.

The Bayesian approach for parameter estimation for ODE modeling has gained attention in later years [14, 22] since it provides an elegant way of addressing the uncertainty associated with the estimated parameters and includes clinical knowledge. The Bayesian approach gives a complete image of the parameter estimation in terms of uncertainty quantification, i.e., posterior mean, credibility intervals and correlations. A Bayesian approach for patient-specific parameters for hemodialysis has been proposed by Bianchi et al. [4] but does not consider the phosphate kinetics.

### 1.2 Contribution

We propose a Bayesian approach for estimating patient-specific parameters for phosphate dynamics during hemodialysis. Moreover, we include the phosphate concentration in the inner compartment as a parameter of the model. We use uncertainty quantification to assess the reliability of our parameter estimates and explore the full parameter space. We address the identifiability of the parameters for the SP, MP and the combination of the two, denoted combined pass (CP). In addition, we also investigate how the parameter estimation can be improved by including relapse measurements and / or measure consecutive sessions.

### 1.3 Outline

Section 2 describes the phosphate kinetics during hemodialysis and introduces the single- and multiple-pass treatments. Section 3 introduces the Bayesian model and describes implementation and sampling diagnostics. In Section 4, we test and validate SP, MP and CP models on data sets and discuss findings from synthetic data which are found in the supplementary materials. Lastly, we conclude the paper in Section 5.

## 2 Hemodialysis modeling

About 85% of the total phosphate in the human body is stored in the bones [15]. We assume that we have an inexhaustible source (bone) that excretes phosphate to the blood, including extracellular fluid. The phosphate transport from source to blood is driven by diffusion. The diffusion process is governed by the diffusion coefficient (permeability) and concentration gradient. The blood compartment is coupled to the dialysate compartment through a semipermeable membrane which generates a flow of phosphate to the dialysate fluid. The flow of phosphate from blood to dialysate is mainly governed by diffusion and to an insignificant degree by a convection process. [17] However, comprehensive investigations have shown that the convective flow has a negligible effect on the model and parameters during the normal range of dialysis treatment, i.e., up to eight hours [2]. Thus, we exclude the convection term from the models. In this paper, we consider three types of models for HD for phosphate clearance in dialysis patients, the conventional SP, MP and the combination CP.

The value of this analysis for clinicians is twofold. Firstly, accurate modeling permits the prediction of phosphate removal during different forms of dialysis, e.g., short and long dialysis or use of filters with standard or high phosphate clearances. Secondly, it is possible to get insight into the underlying physiological causes of phosphate dynamics.

### 2.1 Single-pass dialysis

For the SP treatment, the dialysate is constantly replenished by fresh dialysate such that the phosphate concentration in the outflowing dialysate remains low. Data shows this phosphate concentration to be approximately constant throughout the treatment. SP requires excessive amounts of dialysate for each session. A conceptual diagram of the SP treatment is depicted in Figure 1 that illustrates the removal of phosphate by diffusion.

**Figure 1:**
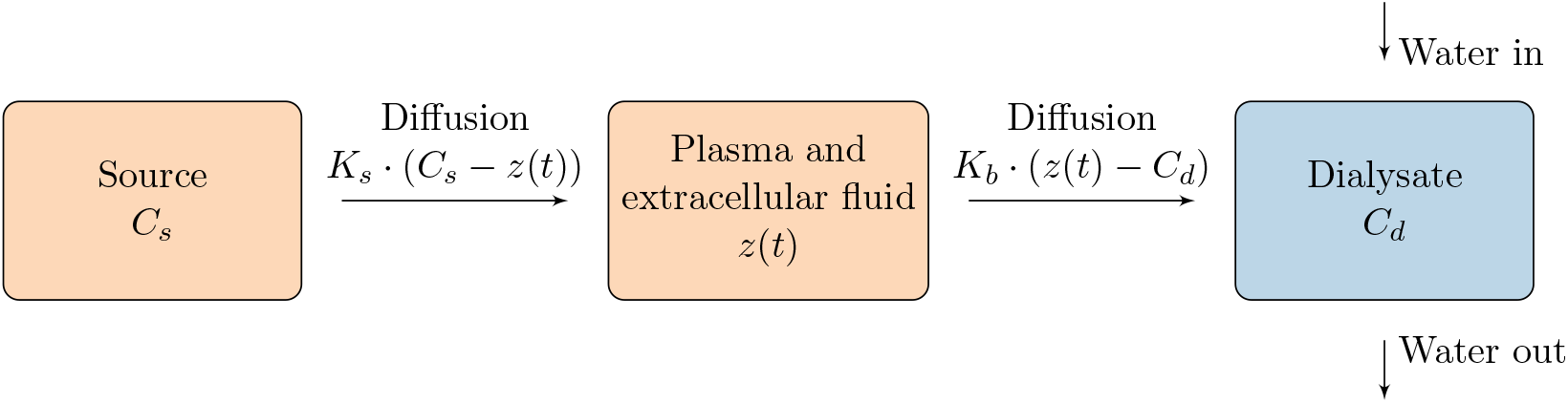
Conceptual diagram for single pass (SP). In SP, blood and dialysate are passed through a filter which initiates a diffusion process that removes toxic substances from the blood (plasma and extracellular fluid). The outflowing dialysate is constantly replenished by fresh dialysate, and the concentration of phosphate in the dialysate is assumed constant.

Agar et al. [1] proposed a simple compartment model for SP consisting of a single linear autonomous ODE,

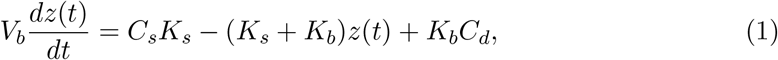

where *z*(*t*) is the concentration of phosphate in the blood compartment at time *t, C*_*s*_ is the constant concentration in the source compartment and *C*_*d*_ is the phosphate concentration in the dialysate assumed to be constant and measurable. *K*_*s*_ and *K*_*b*_ are diffusion rates from source to blood and from blood to dialysate, respectively. Lastly, *V*_*b*_ denotes the blood volume taken as the blood plasma and extracellular volume. For the system to have a unique solution, we equip the ODE with the initial condition *z*(0) = *z*_0_. Notice that the system is not identifiable since *V*_*b*_ can be integrated in the remaining parameters and thus we assume that *V*_*b*_ is known through measurements for single-pass.

The assumption of a constant *C*_*d*_ is not crucial. If we allow the phosphate concentration to be a variable with initial value 0, then we can extend the model by an extra differential equation. This extension results in a fast transient in *C*_*d*_ toward the steady state value given by data shown in Table 1 with at doubling time of approximately 10-15 minutes (see supplementary, Figure S.14). Moreover, such extension does not affect the parameter estimates achieved. Hence we confine ourselves to consider *C*_*d*_ as a constant.

**Table 1:** The mean concentration of phosphate in the dialysate for SP, *C*_*d*_, the mean dialysate volume for MP, *V*_*d*_ and the mean extracellular volume, *V*_*b*_ for both SP and MP.

| Estimate |  | Patient |  |  |  |  |  |  |  |  |  |
| --- | --- | --- | --- | --- | --- | --- | --- | --- | --- | --- | --- |
|  |  | 1 | 2 | 3 | 4 | 5 | 6 | 7 | 8 | 9 | 10 |
| SP | $C_d$ [mmol/L] | 0.16 | 0.12 | 0.11 | 0.16 | 0.18 | 0.21 | 0.14 | 0.09 | 0.14 | 0.09 |
| | $V_b$ [L] | 16.88 | 17.74 | 16.92 | 21.20 | 18.20 | 14.74 | 15.32 | 13.04 | 20.20 | 18.00 |
| MP | $V_b$ [L] | 16.99 | 17.80 | 17.43 | 21.20 | 18.39 | 15.15 | 15.48 | 13.76 | 20.21 | 18.25 |
| | $V_d$ [L] | 22.61 | 23.00 | 26.57 | 31.93 | 28.51 | 20.15 | 23.42 | 14.24 | 28.59 | 23.25 |

### 2.2 Multiple-pass dialysis

Contrary to SP where dialysate is constantly replenished, the dialysate for MP is recirculated, and consequently, the removed substances accumulate in the dialysate fluid over time. A conceptual diagram of the MP treatment is depicted in Figure 2.

**Figure 2:**
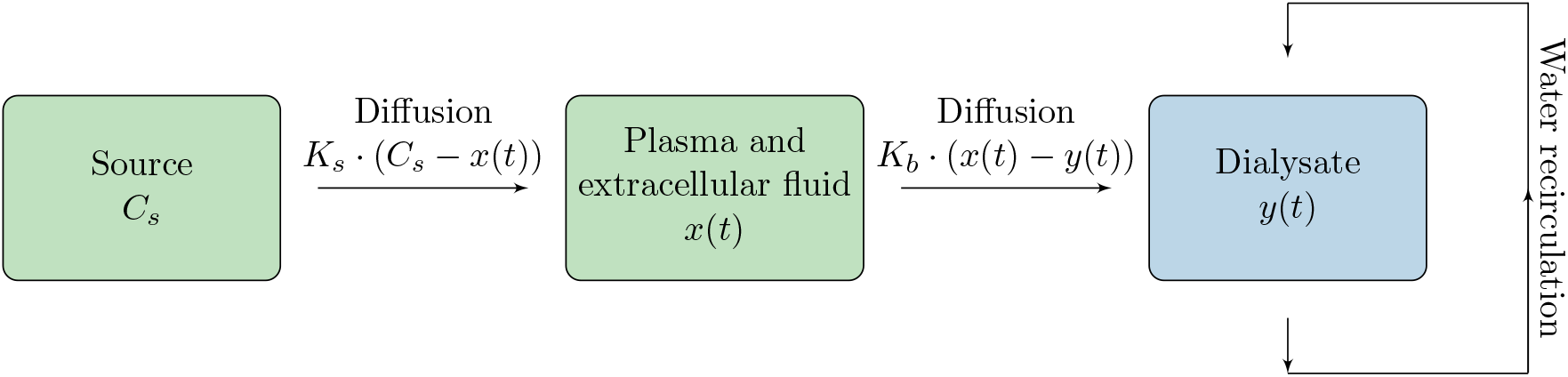
Conceptual diagram for multiple pass (MP) treatment. Like in conventional SP, blood and dialysate is passed through a filter that causes a diffusion process that removes toxic substances from the blood (plasma and extracellular fluid). The dialysate is recirculated and consequently, the removed substances accumulate in the dialysate, i.e., *y*(*t*) changes as a function of time.

MP is less effective than SP due to the accumulation of substances in dialysate. However, MP greatly reduces the amount of dialysate fluid needed for HD treatment, which makes a smaller clinical setting possible. Furthermore, it may ease HD treatment at home and treatment during travels, which can possibly greatly improve the quality of life for renal failure patients. [7, 8, 12]

The MP model can be described by the following system of linear autonomous ODEs,

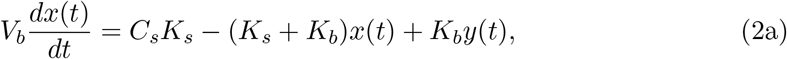

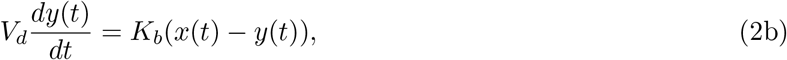

where *x*(*t*) and *y*(*t*) are the time-varying phosphate concentrations for the blood compartment and in the dialysate at time *t*, respectively, and *V*_*d*_ is the volume of the dialysate. The remaining parameters, i.e., *V*_*b*_, *C*_*s*_, *K*_*s*_ and *K*_*b*_, have the same interpretation as for the SP model in (1). The initial conditions are *x*(0) = *x*_0_ and *y*(0) = *y*_0_ corresponding to the phosphate concentration in blood and dialysate at time *t* = 0, respectively. The phosphate concentration in the dialysate at time *t* = 0 is zero, i.e., we assume *y*_0_ = 0 henceforth.

The MP model carries additional information compared to the SP model since the only new parameter, the dialysate volume (*V*_*d*_) is assumed known and can be estimated reliably from available data. Hence, we have an additional equation in the model but the same number of unknown parameters compared to SP. Thus, given sufficient data, the MP model allows for structural identifiability of the parameters due to the addition of (2b) since we cannot simply integrate *V*_*b*_ in the remaining parameters. However, when measurements of *V*_*b*_ are available, we will consider *V*_*b*_ to be known a priori.

### 2.3 Combined-pass dialysis

The parameters for a single patient are shared for the two treatments. Thus, if a patient completes both SP and MP, we can utilize all available information by considering the CP model,

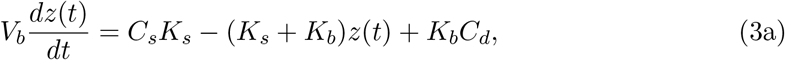

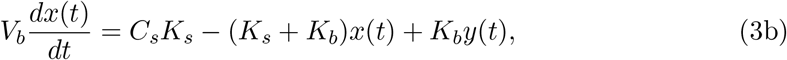

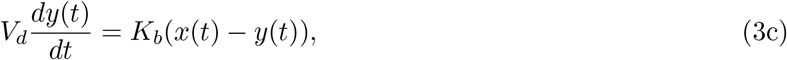

with *z*(0) = *z*_0_, *x*(0) = *x*_0_ and *y*(0) = 0, and the parameters as described for SP. The CP model, just as the MP model, allows for structural identifiability, and potentially even more precise estimation compared to the MP model due to the addition of the SP model.

### 2.4 Clinical data

We consider longitudinal data sets from 10 patients with renal failure that were measured during an SP session and an MP session. The measured phosphate concentrations for SP and MP (*Z, X* and *Y*) are depicted in Figure 3. Measurements were once every hour for a total of four and eight hours for SP and MP, respectively. No measurements were taken in the relapse phase, i.e., after ended treatment.

**Figure 3:**
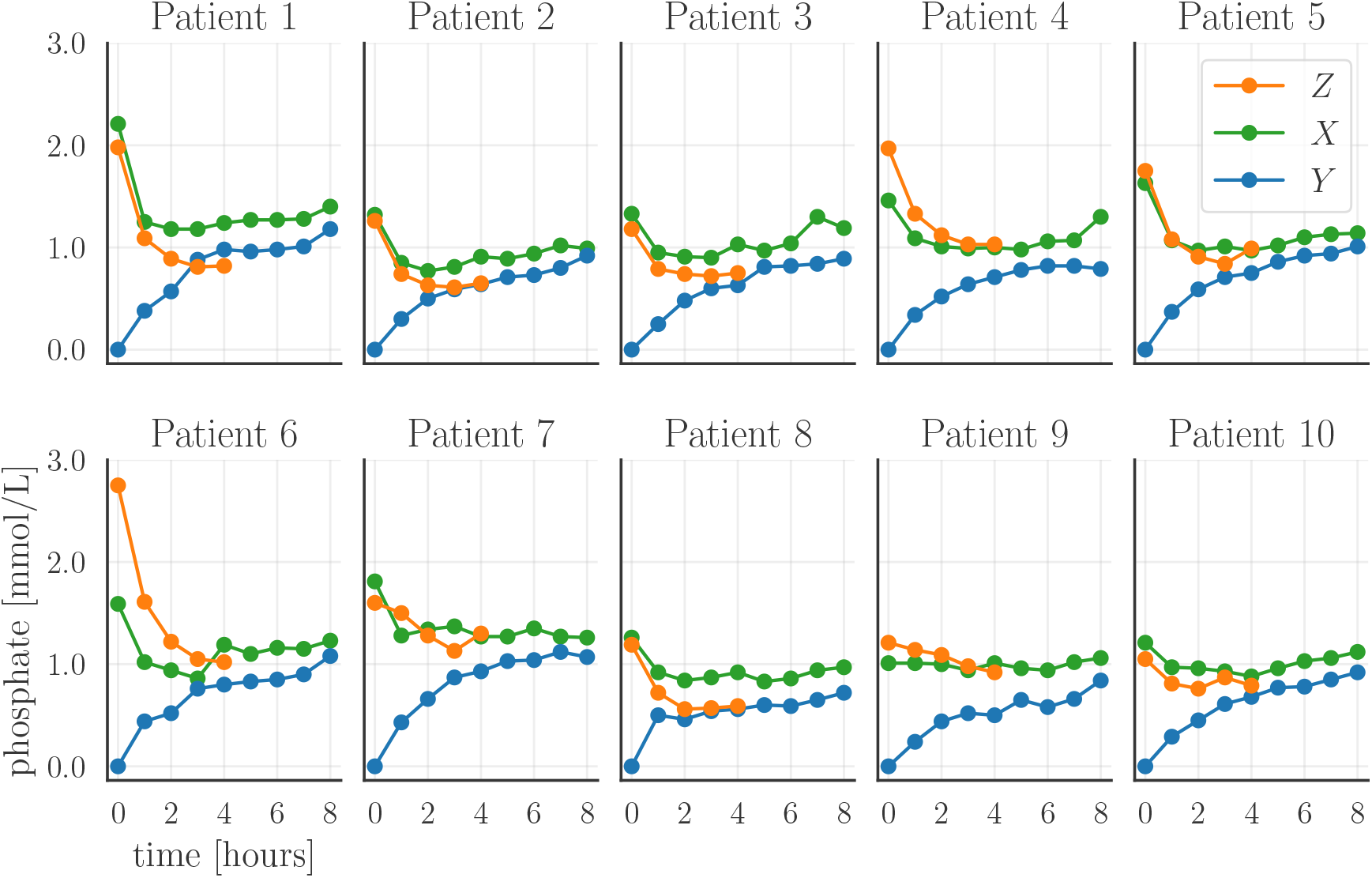
Visualization of the measured phosphate concentrations for SP and MP. The dots represent the measurements, and the full line is the linear interpolation of the measurements. The concentration of phosphate in dialysate in MP is denoted *Y* and the phosphate concentration in the blood is denoted *Z* and *X* for SP and MP, respectively.

Considering the SP measurements (orange dotted line) in Figure 3, we see an exponential-like decay in the measured phosphate concentration after two hours as predicted by (1). Thereafter, phosphate concentration seems to stabilize around a reduced concentration level. For the MP measurements (green and blue dotted lines), we see similar exponential-like decay for the phosphate concentration in agreement with the bi-exponential solution to (2). However, this drop in phosphate concentration is a bit slower for some patients and after two hours it starts to slowly increase due to the accumulation of phosphate in the dialysate. The phosphate concentration in the dialysate increases rapidly in the beginning of the treatment but slows down and approaches an equilibrium with the phosphate concentration in the blood. This behavior is expected according to the model in (2) since the concentration gradient vanishes.

## 3 Bayesian inference

We solve the parameter estimation problem using a Bayesian approach, where we consider the parameters, measurement noise and initial conditions as random variables. In Bayesian inference, we are interested in the posterior probability of the parameters. The posterior probability consists of two components: a prior probability reflecting our knowledge or beliefs about likely parameter values, and a likelihood function that expresses how likely it is to observe the data for a set of parameters. Thus, the posterior allows us to formally include clinical prior knowledge in the model. Moreover, the inclusion of the prior may have a regularizing effect on the parameter estimation problem in the sense that the parameter estimates become less sensitive to measurement noise.

We use uncertainty quantification to assess the reliability of the parameter estimates and the concentrations in terms of posterior statistics, i.e., mean, correlation and 95% credibility intervals (CI). A strength of the uncertainty quantification is that the solution is based on all probable outcomes instead of being solely based on a point estimate [23]. Uncertainty quantification can also be used for model analysis and improvement, e.g., revealing strong correlation or identifying potential measurements that could improve identifiability of the model [26]. Hence, uncertainty quantification is a flexible method to assess how certain we are of the parameter values and parameter-dependent solutions

### 3.1 Likelihood and prior modeling

We describe the Bayesian model for the SP and MP treatments and presume data for the relevant state variables (phosphate concentrations) are measured. Notice that the Bayesian formulation is trivially extended to CP by combining the SP and MP models.

#### 3.3.1 Single-pass formulation

First, we consider the Bayesian formulation for SP. Let *θ* = [*C*_*s*_, *K*_*s*_, *K*_*b*_] denote the vector of unknown parameters and *z*_IC_ denote the initial condition for SP. We assume that *V*_*b*_ and *C*_*d*_ are known to a sufficient degree a priori and do not estimate them based on the model. A justification of this assumption is given in Section 4.

The state variable *z*(*t, θ, z*_IC_) is the solution to (1) and we wish to infer the model parameters *θ* and initial condition *z*_IC_ defining the state variable. Henceforth, we shorten notation such that *z*(*t*) ≡ *z*(*t, θ, z*_IC_).

We assume that the measurement noise is normally distributed such that the state variable, *z*(*t*) is inferred through the Gaussian likelihood function,

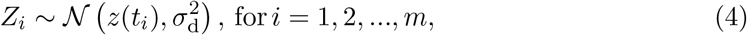

where *Z* ∈ ℝ^*m*^ is a vector with the measurement of the phosphate concentration in the blood at time *t* = *t*_*i*_. The parameter 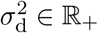 is a hyperparameter describing the variance of the measurement noise. The hyperparameter 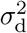 is not known a priori. Thus, we infer 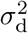 as a parameter of the model and assign an inverse gamma prior [23]. We enforce non-negativity on the likelihood function by truncating it at 0 since the phosphate concentrations are non-negative.

We consider the initial condition *z*_IC_ to have mean equal to the phosphate concentration at time *t* = 0 and variance 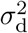 equal to the measurement error, i.e.,

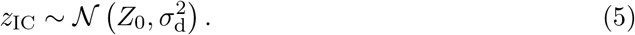

This choice of prior for the initial condition can be interpreted as the initial measurement following the same measurement model as the measurements for time *t >* 0, i.e., we do not assume that the first measurement is more accurately measured than the subsequent ones.

We model the prior of the unknown parameters *θ* by the Gaussian distribution,

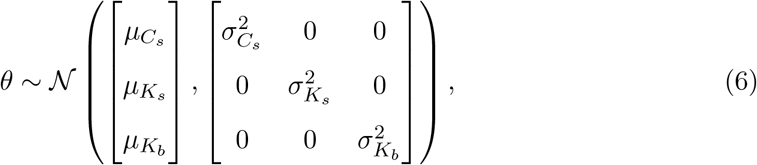

where 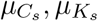 and 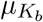 represent the prior clinical knowledge, i.e., our prior belief about most likely parameter values 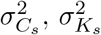 and 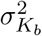 are the variances for *C*_*s*_, *K*_*s*_ and *K*_*b*_, respectively. As with the likelihood function, we impose constraints such that we only consider the parameters in a physiologically meaningful range.

As commonly done, we assume that at the start of the dialysis, i.e., *t* = 0, the patient’s phosphate concentration is approximately in a steady state, i.e., we assume that *Z*_0_ is close to *C*_*s*_ and we choose 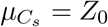. The steady state assumption follows from (1) where *K*_*b*_ = 0 when the patient is not receiving dialysis treatment.

In previous publications [1, 2, 5] *C*_*s*_ is fixed to the value of the initial phosphate measurement. However, the data from Agar et al. [1] show large uncertainty for the first measurement point. Our choice of prior allows *C*_*s*_ to deviate from the initial measurement of the phosphate concentration and thereby our model is not oblivious to measurement errors for the initial phosphate measurement.

We base our values for 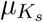 and 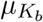 on literature and we choose 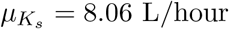 and 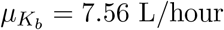 [5].

We initially considered 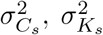 and 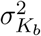 to be parameters of the model. However, preliminary results showed that it greatly decreased the stability of the results. Thus, we choose 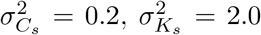 and 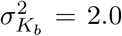 based on visual inspection of the prior to incorporate adequate uncertainty about the prior mean.

#### 3.1.2 Multiple-pass formulation

The main difference between the MP formulation and the SP formulation is the inclusion of an additional state variable through equation 2b. Hence, the likelihood function for MP is

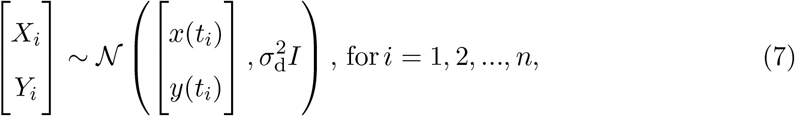

where *X* ∈ ℝ^*n*^ and *Y* ∈ ℝ^*n*^ are vectors with the measurements of the phosphate concentration in the blood and dialysate at time *t* = *t*_*i*_, respectively, and *I* is the 2 × 2 identity matrix. The initial condition for the phosphate concentration in the dialysate is set to zero, i.e., *y*_*IC*_ = 0, and the initial condition for the phosphate concentration in the blood is assigned a prior with mean *X*_0_ and variance equal to the measurement variance, i.e.,

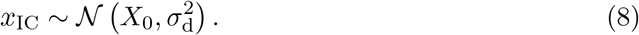

Lastly, we choose the prior for the parameters *θ* to be (6) with the exception that 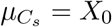.

### 3.2 Implementation and diagnostics

We use sampling-based techniques to approximate the posterior [26]. Markov Chain Monte Carlo (MCMC) is a sampling technique that generates a Markov chain of samples that converges to the posterior distribution of the parameters [21]. Hence, we can compute posterior statistics, i.e., mean, 95% CI and correlation from the Markov chain.

The simple MCMC techniques such as random walk Metropolis Hastings and the Gibbs sampler are plagued by inefficient exploration of the parameter space via random walks and are highly sensitive to correlated parameters. Hamiltonian Monte Carlo (HMC) is an MCMC method that avoids random walk behavior by taking a series of first-order gradient informed steps in the simulation and explores the parameter space well even in the case of correlated parameters. The performance of the HMC sampler is highly sensitive to the choice of user-specified parameters. However, the No-U-Turn Sampler (NUTS) is an HMC method where the user-specified parameters are automatically estimated. [13] We use Runge-Kutta 45 (RK45) to solve the ODE system [19] and the PySTAN implementation of NUTS [9] with default choice for all associated parameters to compute the samples that approximate the posterior distribution.

For each simulation, we generate four sample chains from random initializations, and we consider the potential scale reduction statistic, the so-called 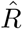 value for sampling diagnostics [10]. The 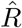 value measures the ratio of the average variance of samples within each chain to the variance of the pooled samples across chains, and if all chains are at equilibrium, then the 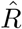 value will be one.

## 4 Results

In this section, we consider two data sets for dialysis patients during hemodialysis. For each patient, we generate 4000 samples and visualize the results in terms of posterior mean and 95% CIs for the estimated parameters and phosphate concentrations during and after hemodialysis. In addition, we also visualize the pairwise correlation for the parameters by scatter plots of the samples and compute the relative standard deviation. All presented results returned an 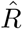 value of one, indicating convergence of the sample chains. In addition, we visually inspected the sample chains, which appeared well mixed. Tables with estimated posterior means, 95% CIs and relative standard deviations are found in Appendix B, and RMSE is listed in Table 2.

**Table 2:** Computed RMSE for Figure 4, 7 and 8. We compute RMSE by the formula 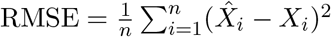 where 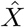 and *X* are the estimated and measured phosphate concentrations, respectively.

| Patient | SP | MP | CP |
| --- | --- | --- | --- |
| 1 | $2.14 \cdot 10^{-8}$ | $2.08 \cdot 10^{-4}$ | $1.67 \cdot 10^{-3}$ |
| 2 | $3.23 \cdot 10^{-6}$ | $5.06 \cdot 10^{-6}$ | $2.12 \cdot 10^{-4}$ |
| 3 | $1.69 \cdot 10^{-6}$ | $1.95 \cdot 10^{-5}$ | $2.59 \cdot 10^{-4}$ |
| 4 | $7.99 \cdot 10^{-9}$ | $1.79 \cdot 10^{-5}$ | $9.95 \cdot 10^{-4}$ |
| 5 | $3.79 \cdot 10^{-5}$ | $1.85 \cdot 10^{-7}$ | $1.11 \cdot 10^{-3}$ |
| 6 | $6.36 \cdot 10^{-9}$ | $3.99 \cdot 10^{-5}$ | $1.67 \cdot 10^{-3}$ |
| 7 | $4.88 \cdot 10^{-5}$ | $7.56 \cdot 10^{-5}$ | $1.25 \cdot 10^{-2}$ |
| 8 | $2.22 \cdot 10^{-7}$ | $4.27 \cdot 10^{-5}$ | $5.63 \cdot 10^{-4}$ |
| 9 | $2.87 \cdot 10^{-6}$ | $4.84 \cdot 10^{-5}$ | $8.18 \cdot 10^{-4}$ |
| 10 | $9.07 \cdot 10^{-6}$ | $9.41 \cdot 10^{-7}$ | $2.46 \cdot 10^{-4}$ |

We have also investigated the models using synthetic data to confirm the findings of the results with real data. These synthetic experiments can be found in the supplementary. Here we present the results obtained by the Bayesian model described in Section 3 for the data depicted in Figure 3.

### 4.1 Single pass and Multiple pass

First, we consider the hemodialysis data for the ten patients shown in Figure 3. Beside phosphate concentrations in the blood and dialysate depicted, we have hourly measurements of the phosphate concentration in the dialysate (*C*_*d*_) for SP, the volume of the blood compartment (*V*_*b*_) for both SP and MP, and the dialysate volume (*V*_*d*_) for MP. *C*_*d*_ was measured when exiting the dialysate compartment after initializing the dialysis process. We assume that the concentration of phosphate in the dialysis for SP is constant as suggested by data, and for each patient, we compute *C*_*d*_ as the spatial average of the concentration of phosphate from inlet to outlet of the dialysis machine. Table 1 lists *C*_*d*_, *V*_*d*_ and *V*_*b*_ estimated directly from available data and Figure A.1 and Figure A.2 in Appendix A provide exploratory statistics of the corresponding data.

#### 4.1.1 Estimation

The estimated phosphate concentrations obtained for SP are depicted in Figure 4 along with the predicted relapse. The solid line represents the posterior mean, the full circles are data points and the transparent region indicates the 95% CI i.e., the region that contains 95% of the samples. Considering the estimated phosphate concentrations for SP, we see that the sampler has computed a decent fit in terms RMSE in Table 2 and posterior mean with a narrow 95% CI for the treatment phase. However, there is a large 95% CI for the relapse phase.

**Figure 4:**
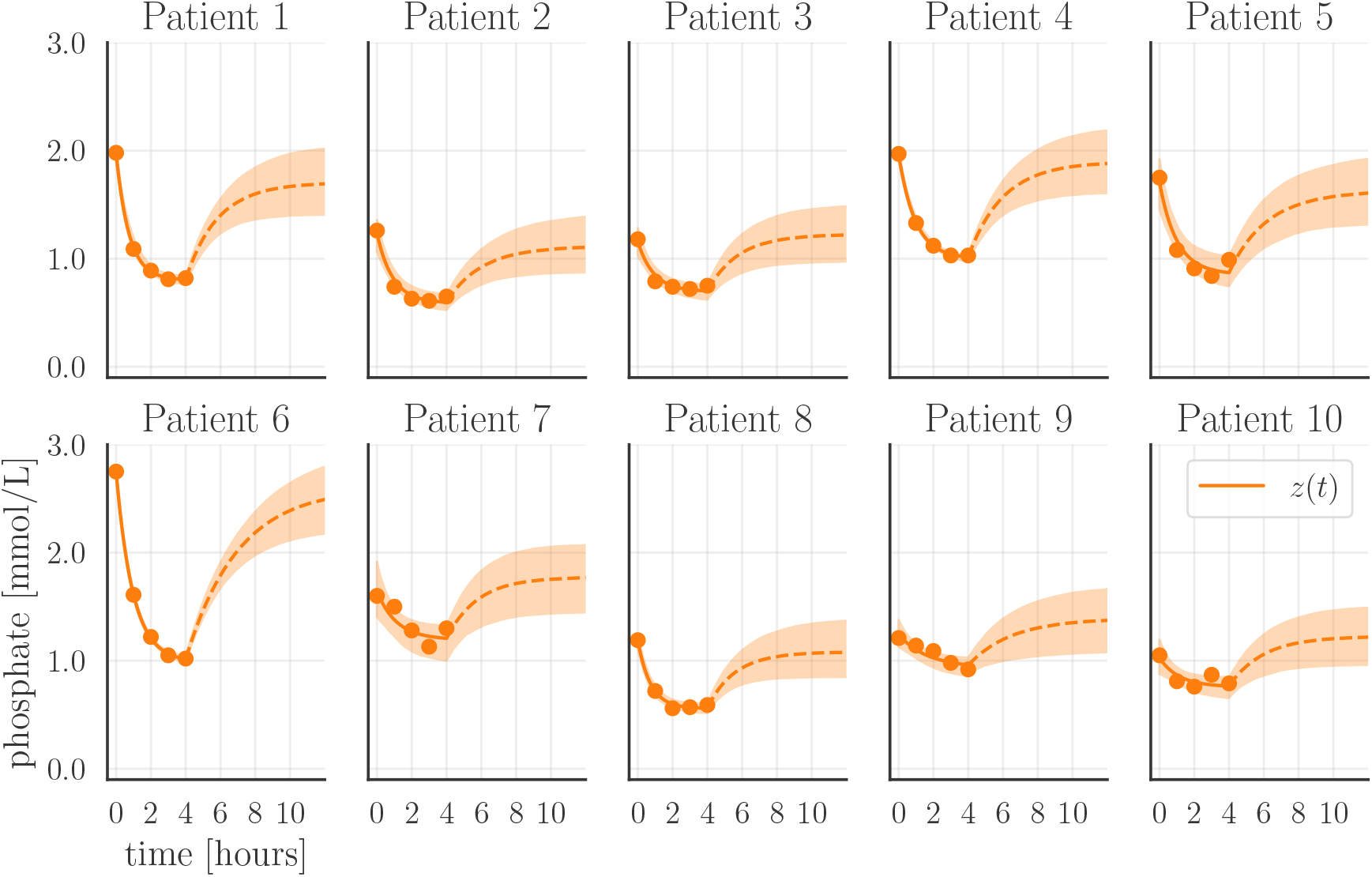
Estimated treatment and relapse for SP. The full lines are the posterior mean of the samples whereas the transparent regions represent the 95% CI. The full circles are measurements, and the dashed line is the posterior mean of the estimated relapse. For RMSE, see Table 2.

The corresponding parameter estimates with 95% CI for SP are visualized in Figure 5 and listed in Table B.1 where the average relative standard deviation is 10.3%, 18.4% and 18.6% for *C*_*s*_, *K*_*s*_ and *K*_*b*_, respectively. The full posterior density for the parameters for patient 2 is shown in Figure 6. We have chosen to only include a correlation plot for patient 2 in this section since it shows the general trend of the estimated parameters. The correlation plots for the remaining patients are found in Figure C.1-C.9 in Appendix C.

**Figure 5:**
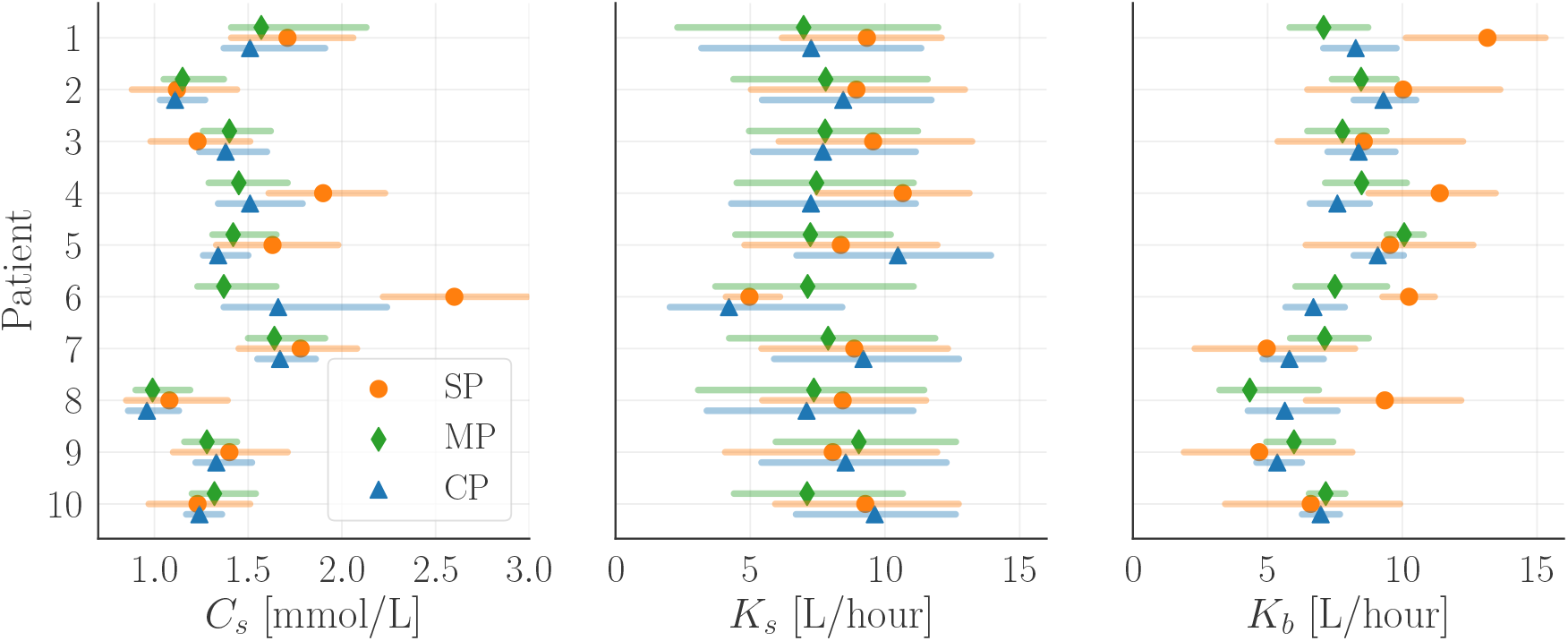
Visualization of the parameter estimates. The dots, diamonds and triangles represent the posterior mean for SP, MP and CP, respectively. The transparent region is the 95% CI. The full posterior of the parameters for patient 2 is shown in 6 and for the remaining patients in Figure C.1-C.9 in Appendix C.

**Figure 6:**
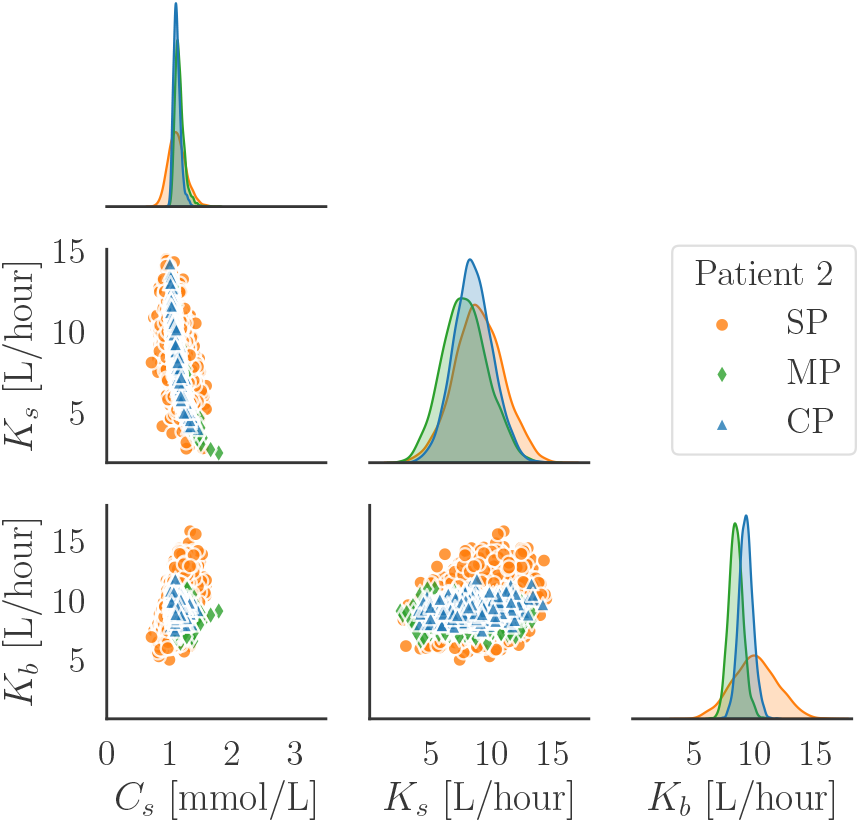
Plot of the posterior density and correlation of the parameters estimated for patient 2 for SP, MP and CP. The density plots show the posterior density functions, and the scatter plots show the posterior samples.

Figure 7 shows the estimated phosphate concentrations and predicted relapse phase for MP. Figure 7 and Table 2 show that the parameter estimation has found a satisfying fit both visually and in terms of RMSE for MP as for SP. However, the width of the 95% CIs is smaller for the relapse phase. The reduced uncertainty in the relapse can be explained by the reduced 95% CI for *C*_*s*_ in MP compared to SP which is shown in Figure 5 and Figure 6 and quantified by the decreased average standard deviation of 7.3% in Table B.2, i.e., a reduction of 3%.

**Figure 7:**
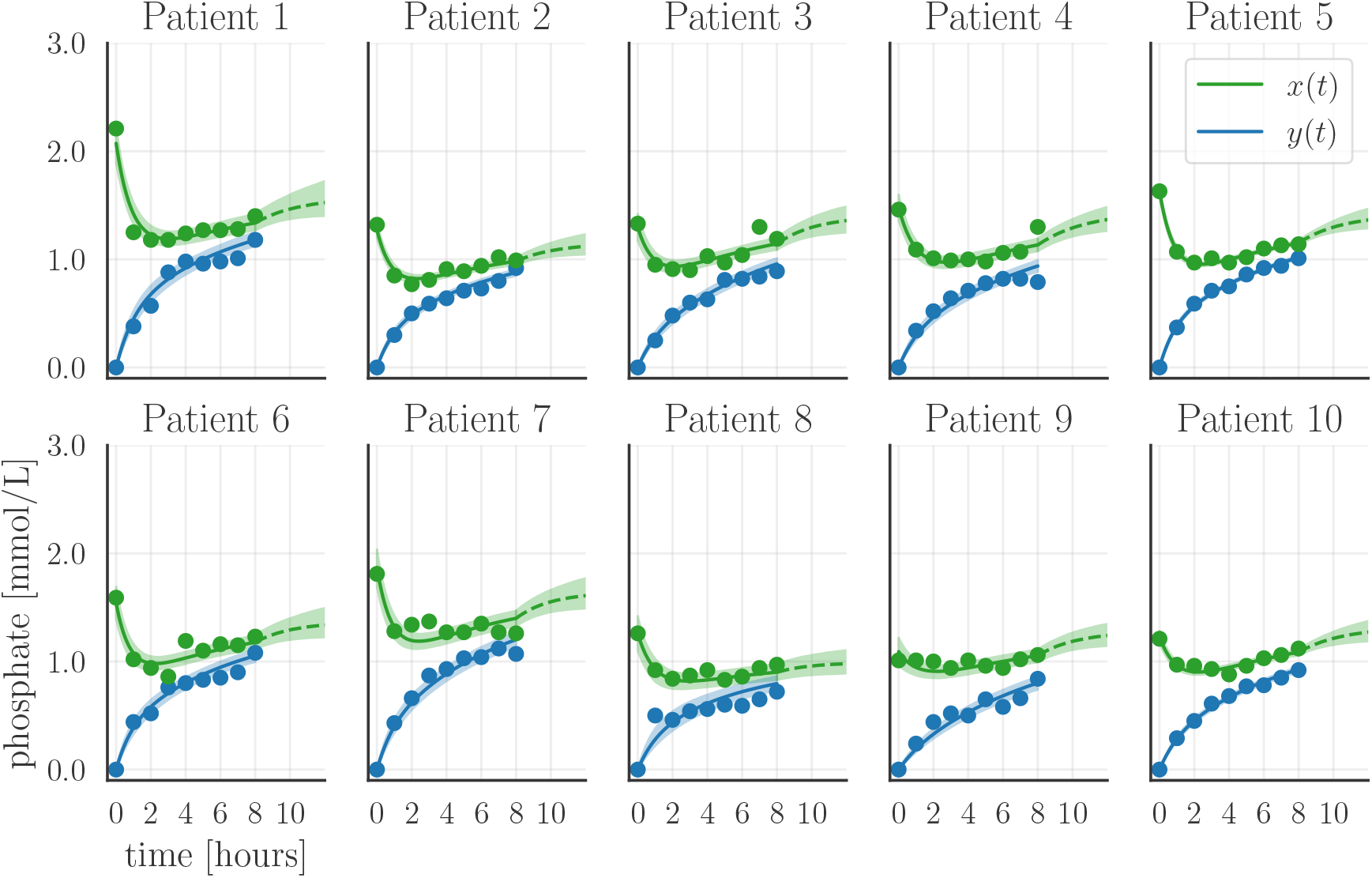
Estimated treatment and relapse for MP. The full lines are the posterior mean of the samples whereas the transparent regions represent the 95% CI. The full circles are measurements, and the dashed line is the posterior mean of the estimated relapse. For RMSE, see Table 2.

Moreover, Figure 5 shows a great reduction in the uncertainty about *K*_*b*_ as expected from the addition of equation (2b) with a relative standard deviation of 9.6%, i.e., a reduction of 9% compared to SP. However, the uncertainty about *K*_*s*_ remains largely unaffected by the additional knowledge utilized by the MP model and the uncertainty actually increases on average with an average relative standard deviation of 24.8%.

Considering the CP results in Figure 8, we see that CP finds a unified set of parameters that describe the SP and MP sessions for each patient. Moreover, the CP estimates a satisfying fit both visually and in terms of RMSE in Table 2. The parameter estimates are very similar to the ones obtained by MP as seen in Figure 5 except for patient 6 and with only a slight reduction compared to MP in average relative standard deviation, 6.9%, 22.9% and 8.2% for *C*_*s*_, *K*_*s*_ and *K*_*b*_, respectively. A possible explanation for the difference in *K*_*s*_ for patient 6 is the large difference in initial measured phosphate concentration, indicating that steady state had not been reached before treatment onset.

**Figure 8:**
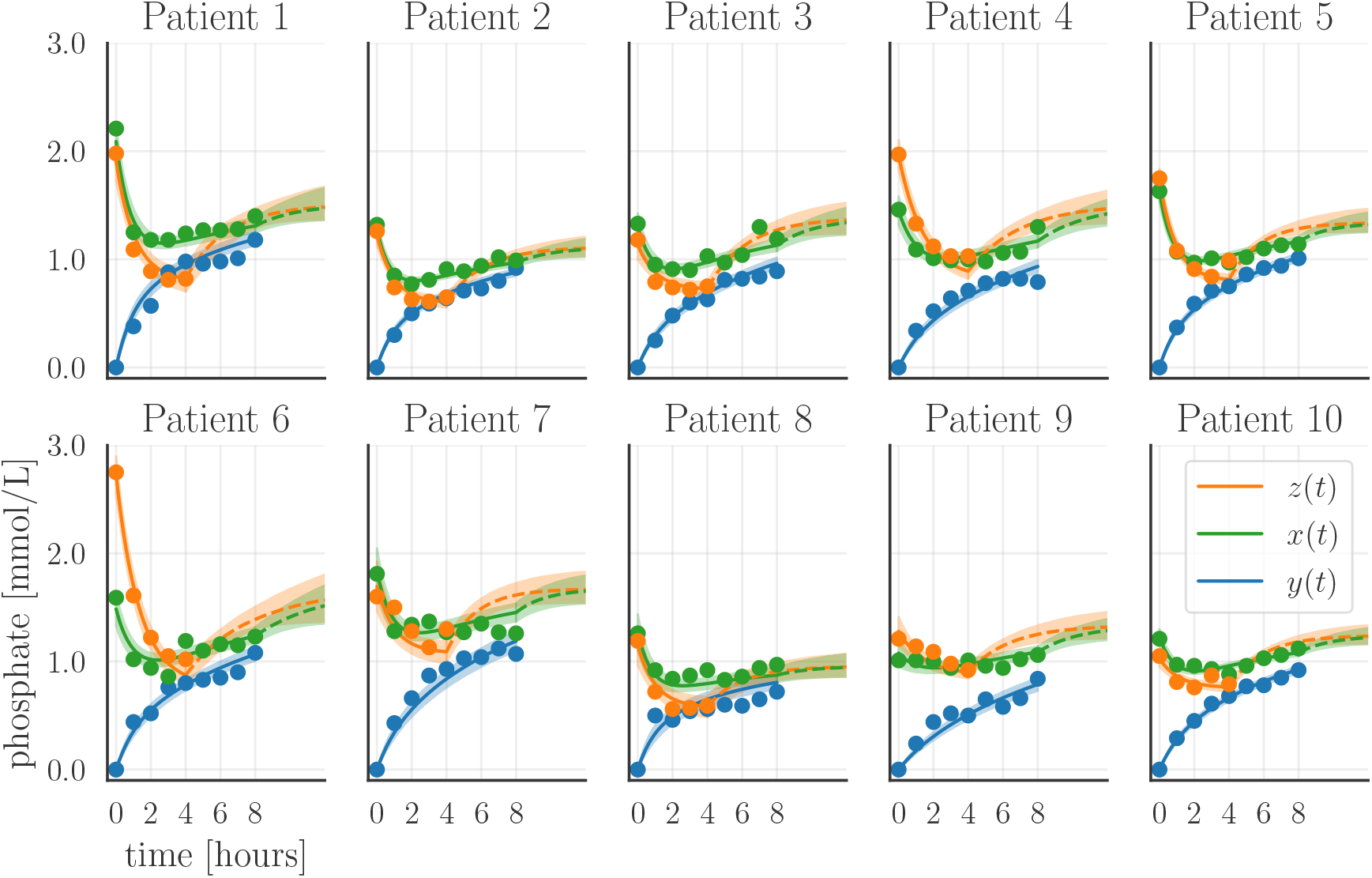
Estimated treatment and relapse for CP. The full lines are the posterior mean of the samples whereas the transparent regions represent the 95% CI. The full circles are measurements, and the dashed line is the posterior mean of the estimated relapse. For RMSE, see Table 2.

The synthetic results in Figure S.2 and S.3 in the supplementary materials show that with fixed *C*_*s*_ and a uniform prior on *K*_*s*_ and *K*_*b*_ (mimicking the parameter estimation in [1, 5, 2]), we have a very limited identifiability of *K*_*s*_ and *K*_*b*_ for SP, whereas MP and CP recover values very close to the true parameters with significantly lower uncertainty. In addition, the parameter estimates for *K*_*s*_ and *K*_*b*_ were highly correlated and this correlation was significantly reduced by MP and CP. We also considered the full Bayesian model with priors on the synthetic data and the results are depicted in Figure S.5 and S.6 in the supplementary materials. The results showed that MP and CP in general came closer to the true parameters with smaller 95% CI and showed similar results in terms of relative standard deviation.

In summary, the uncertainty associated with the SP results is reduced significantly by using the Bayesian model with priors compared to the standard parameter estimation without the clinical knowledge incorporated. For the Bayesian models, we see that MP and CP are superior to SP in estimating patient-specific parameters *C*_*s*_ and *K*_*b*_, but that the gain of considering CP compared to MP is limited. However, we see that the uncertainty about *K*_*s*_ is large even when using all available data with the CP model. These findings are further supported by the synthetic results in the supplementary materials, where the estimates obtained by MP and CP are closer to the true parameter value and with less uncertainty. Thus, based on the estimation results, it seems that the SP data without relapse data or consecutive sessions are not sufficient for estimating the parameters reliably.

### 4.2 Consecutive SP sessions

Debowska et al. [5] present a data set consisting of 25 patients that were examined during three consecutive SP sessions of a one-week dialysis treatment cycle. They present the data as the average of the measurement for the 25 patients and we have read off the data from the figures. Measurements were obtained hourly for a total duration of four hours with the addition of a measurement 45 minutes after ended treatment, i.e., we have five SP measurements and a relapse measurement for each of the three consecutive SP sessions. We choose *V*_*b*_ = 20 and *C*_*d*_ = 0.

#### 4.2.1 Simulations and estimates

The aim of this subsection is to investigate the improvement of information obtained by including relapse measurement and / or consecutive sessions in the SP model. We investigate the four following scenarios, No Relapse (NR) where we consider the first SP treatment only, Partial Relapse (PR) with the first SP treatment with a measured relapse point, Full Relapse (FR) where we consider the first SP treatment with relapse point and the first measured data point of the second SP, and Full Three Relapse (FTR) where we include the data from all three SP consecutive sessions.

The results for the four scenarios are depicted in Figure 9. The measurements included in each parameter estimation are marked with colored dots, whereas the measurements not included in the model estimation are marked with black open circles. The posterior statistics for the parameters are shown in Figure 10 and listed in Table B.4. Correlation of the parameters is shown in Figure 11.

**Figure 9:**
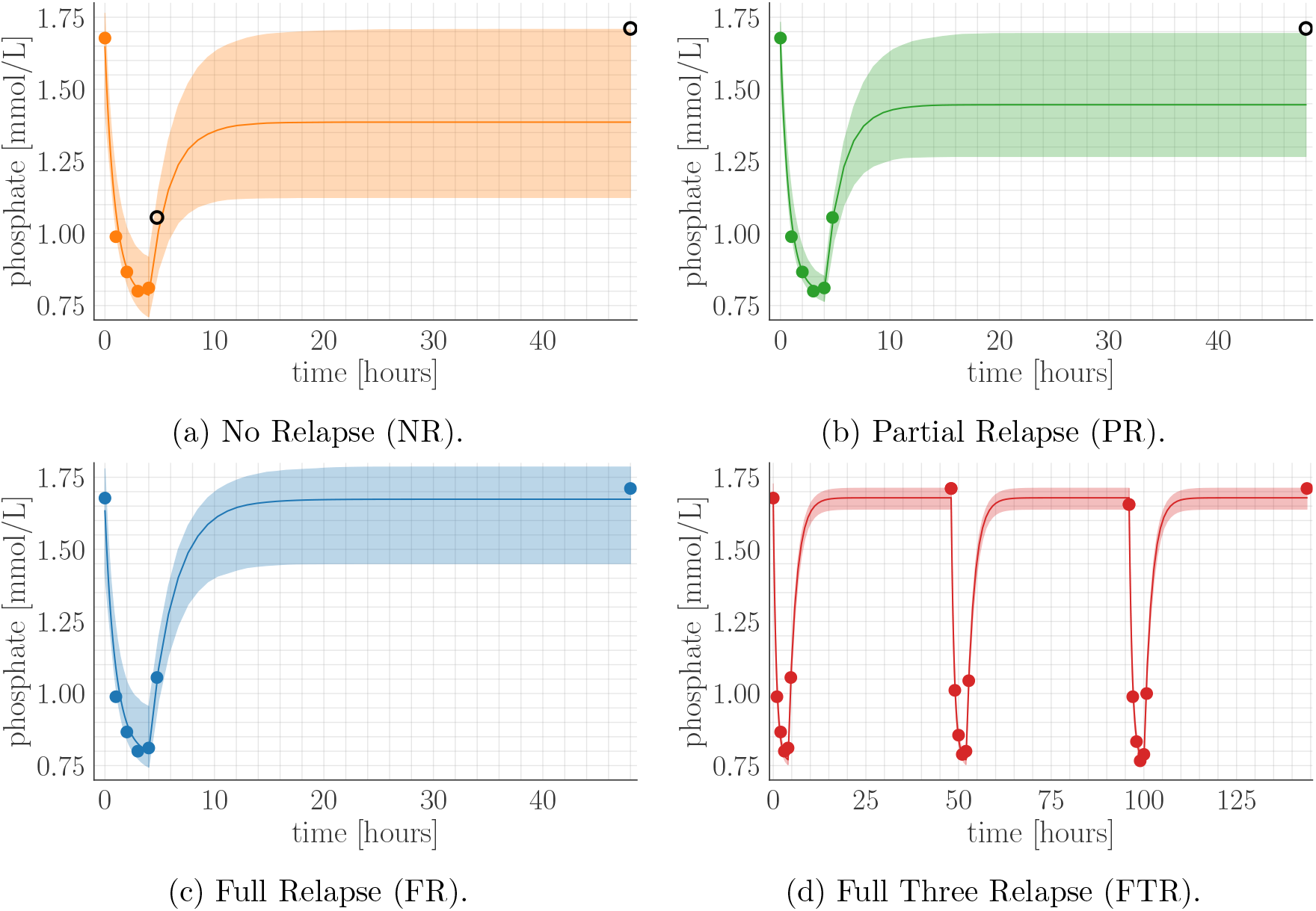
Four simulations for the relapse data. (a) first SP with no relapse data (NR), (b) first SP with a single relapse data point after 4.75 hours (PR), (c) first SP with full relapse data, i.e., after 4.75 and 48 hours (FR). Lastly (d) shows the fit when including all three consecutive SP treatments (FTR). The measurements are shown with colored circles. The open black circles in (a) and (b) indicate that the measurements are not used for estimation. RMSE is NR =6.18·10^−6^, PR=1.19·10^−6^, FR=1.07·10^−5^ and FTR=9.36·10^−7^, respectively.

**Figure 10:**
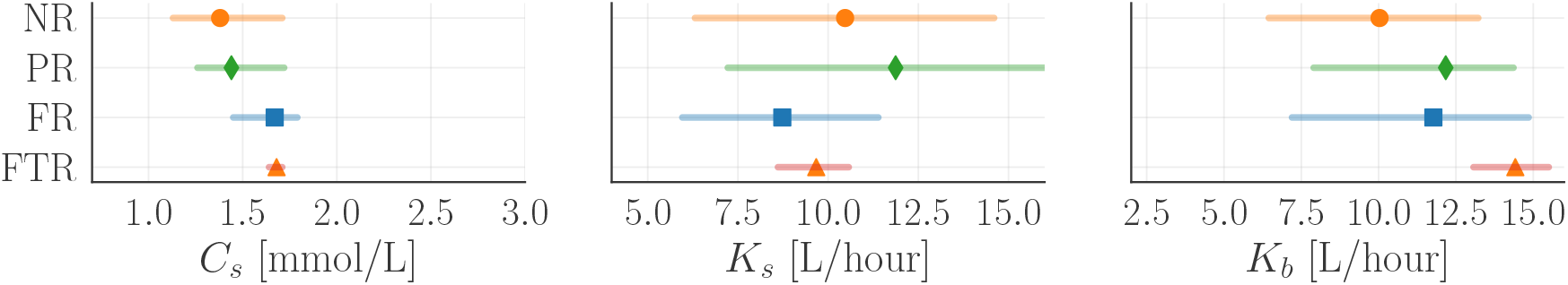
Posterior mean and 95% CI for the parameter estimates for the four scenarios, No relapse (NR), Partial relapse (PR), Full relapse (FR) and full-three relapse (FTR). The figure shows that the uncertainty about the parameter estimates decreases as the number of measurements increases.

**Figure 11:**
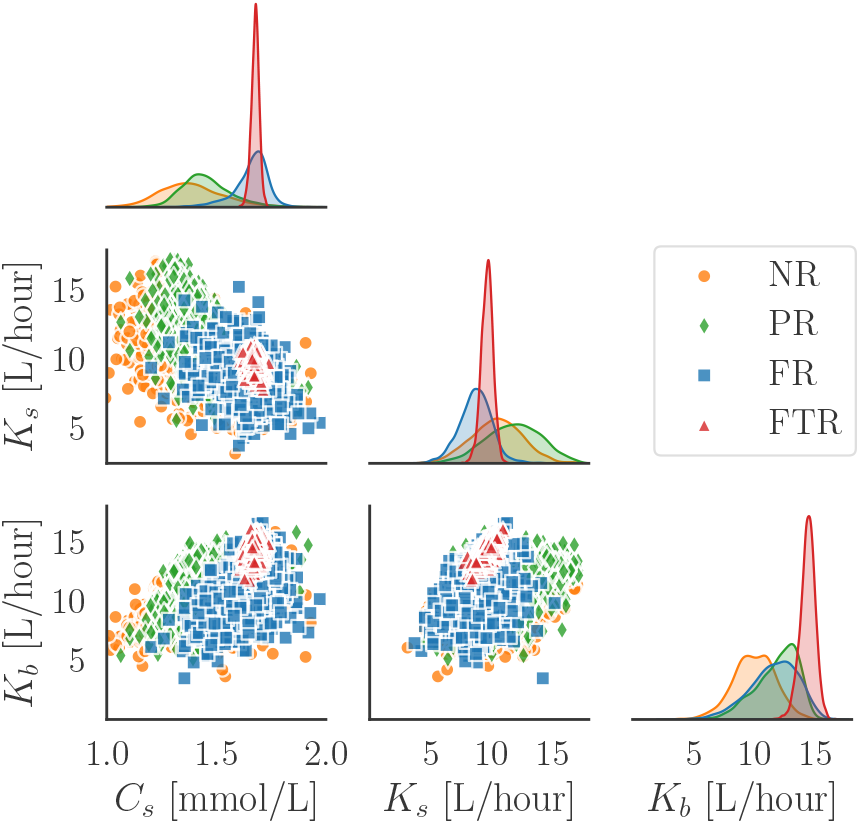
Plot of the posterior density and correlation for the parameter estimates for the four scenarios, No relapse (NR), Partial relapse (PR), Full relapse (FR) and full-three relapse (FTR). The density plots show the posterior density functions, and the scatter plots show the posterior samples. The uncertainty associated with the posterior mean of the parameters decreases as more information is included in terms of relapse measurement and /or consecutive sessions.

Figure 9a shows estimation without relapse measurement for a single SP session, the phosphate concentration has a quite large 95% CI and undershoots the relapse. If we consider the uncertainty in the correlation plot for the parameters in Figure 11 and Figure 10, we see a large 95% CI for the parameter estimates and relative standard deviation in Table B.4 which is similar to the uncertainty associated with the estimate for the SP estimation in Section 4.1.

A model estimation including the measured relapse 45 minutes after ended treatment is depicted in Figure 9b. The 95% CIs for the phosphate concentration is slightly reduced, but the 95% CIs for the parameters have barely changed as seen in Figure 10, Figure 11 and Table B.4. Hence, including a measurement after 45 minutes relapse has limited effect on the uncertainty of the parameter estimates. This can also be seen by considering the correlation plot in Figure 11, where the width of the distribution is only slightly changed. It is noteworthy that the addition of the relapse point has such limited effect on the estimation. However, this limited effect is due to the very rapid dynamics in the initial relapse phase. The initial relapse is not very sensitive to small changes, whereas a relapse point measured later e.g., after two hours, will have a larger effect on the estimation process due to the slower change in the concentration.

Considering the full relapse in Figure 9c, we see the effect of having a relapse measurement several hours after ended treatment. The estimated steady state for the phosphate concentration has an increased posterior mean and narrower 95% CI compared to Figure 9a and 9b. This increase is explained by the increase for *C*_*s*_ which can be seen in Figure 10 and Figure 11. There is also a slight narrowing of the 95% CI for *K*_*s*_ whereas the effect on *K*_*b*_ is limited as the relative standard deviation actually increases from 16% to 18% compared to the partial relapse. Hence, including relapse measurements has limited effect on the identifiability of *K*_*b*_, but reduces the uncertainty associated with the estimates for *C*_*s*_ and *K*_*s*_. This observation is expected based on the model (1), since we have *K*_*b*_ = 0 in the relapse phase.

Lastly, if we have three consecutive SP treatments for the same patient, we can reduce the uncertainty even further, as shown in Figure 9d. The three consecutive SP treatments carry significant information since the repetition makes the estimates less sensitive to fluctuations in the data, which can also be seen in Figure 10 and Figure 11. Considering the relative standard deviation for *K*_*s*_ in Table B.4, we find that it decreases from 20% to 5 % by considering the consecutive sessions compared to a single session. However, even in the case of a single session, our Bayesian approach has significantly smaller relative standard deviation compared to the estimates found by Debowska et al. [5] and Agar et al. [1], who report a relative standard deviation of 79% and 47%, respectively. Even for *K*_*b*_, we see a significant narrowing of the 95% CI. Thus, measuring consecutive sessions greatly increases the identifiability of all three model parameters as the relative standard deviation decreases significantly for all three parameter estimates, as seen in Table B.4.

We also investigated the effect of including relapse measurements for the synthetic data for SP, MP and CP. The results including relapse measurements are shown in Figure S.8-S.11 and results for two consecutive sessions are shown in Figure S.12 and S.13. Here we found that the consecutive sessions were more effective than relapse measurements to reduce the uncertainty of the parameters which aligns with the findings in Figure 9. In general for the synthetic data, we found that MP compared to SP had less uncertainty and came closer to the true parameter values.

## 5 Conclusion

Phosphate clearance with hemodialysis is crucial for patients with renal failure since abnormal levels of phosphate are associated with increased vascular calcification and mortality. We propose a Bayesian approach to parameter estimation for patients undergoing hemodialysis treatments (SP, MP and CP). The Bayesian approach allows us to formally include clinical knowledge in the model and to use uncertainty quantification to assess how reliably we can estimate the three model parameters: phosphate concentration in the bones, phosphate clearance from bone to blood and from blood to dialysate.

We validated and tested our Bayesian model on two data sets for patients with renal failure. The results showed that the uncertainty for the parameter estimates is greatly reduced by considering MP and CP compared to SP. However, for the parameter governing the diffusion rate between bone phosphate and blood, the uncertainty remained unchanged. We also investigated the impact of including relapse data and consecutive treatments. The results showed that including an early relapse measurement (after 45 minutes) had little effect on the estimation process if not combined with a measurement in the later relapse phase. The relapse measurements taken more than 45 minutes after ended treatment had significant impact on the reliability of the model parameters. Moreover, the results showed that we can reduce the relative standard deviation for the phosphate clearance from blood to bone from 20% to 5% by including consecutive sessions in the estimation process compared to estimation based on a single session.

Numerical results on synthetic data confirmed the findings obtained from the real data, and showed that the parameters were poorly identified for SP if no prior information was included. The uncertainty of the estimates greatly decreased when using the Bayesian model incorporating clinical knowledge, and the MP model generally was closer to the true parameter values of the model. Compared to existing parameter estimates of the phosphate clearance from bone to blood, our Bayesian model can estimate a parameter associated with significantly lower uncertainty for both SP and MP.

## Acknowledgments

This work was supported by The Villum Foundation (grant no. 25893).

## A Summary statistics

**Figure A.1:**
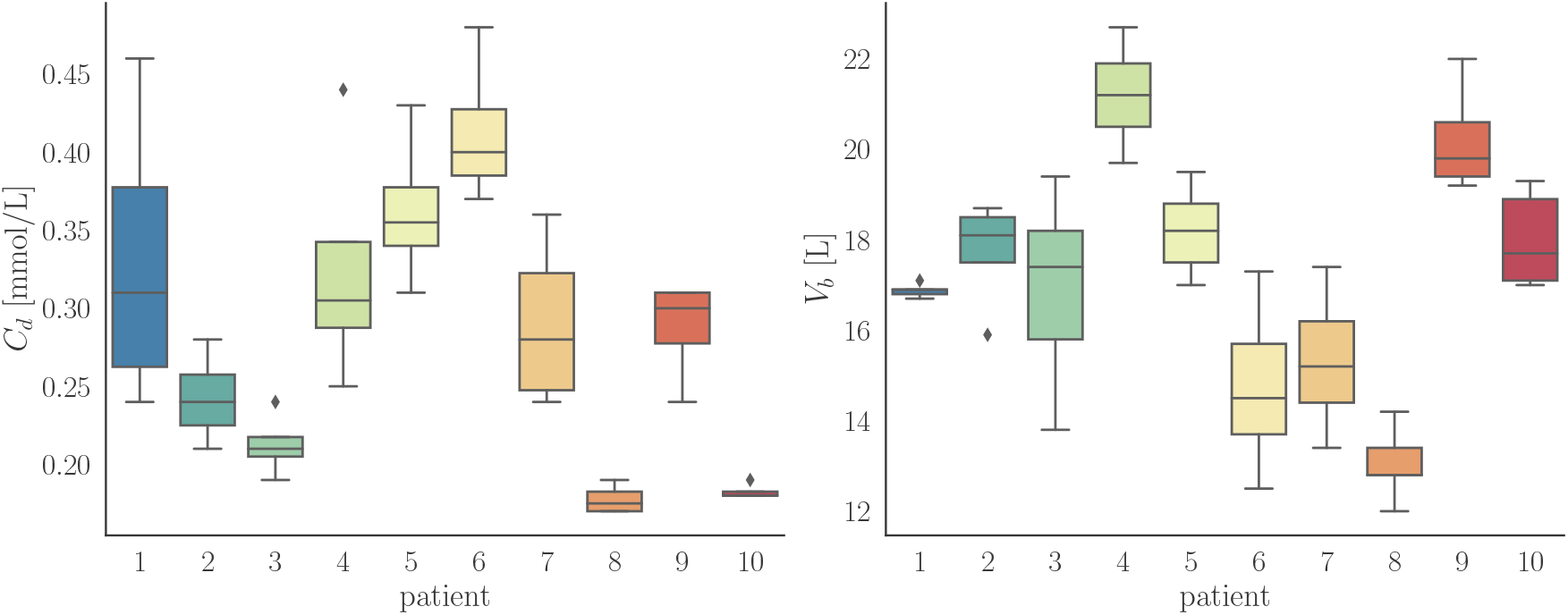
Boxplots of the measured parameters *C*_*d*_ and *V*_*b*_ for the SP sessions in Figure 3.

**Figure A.2:**
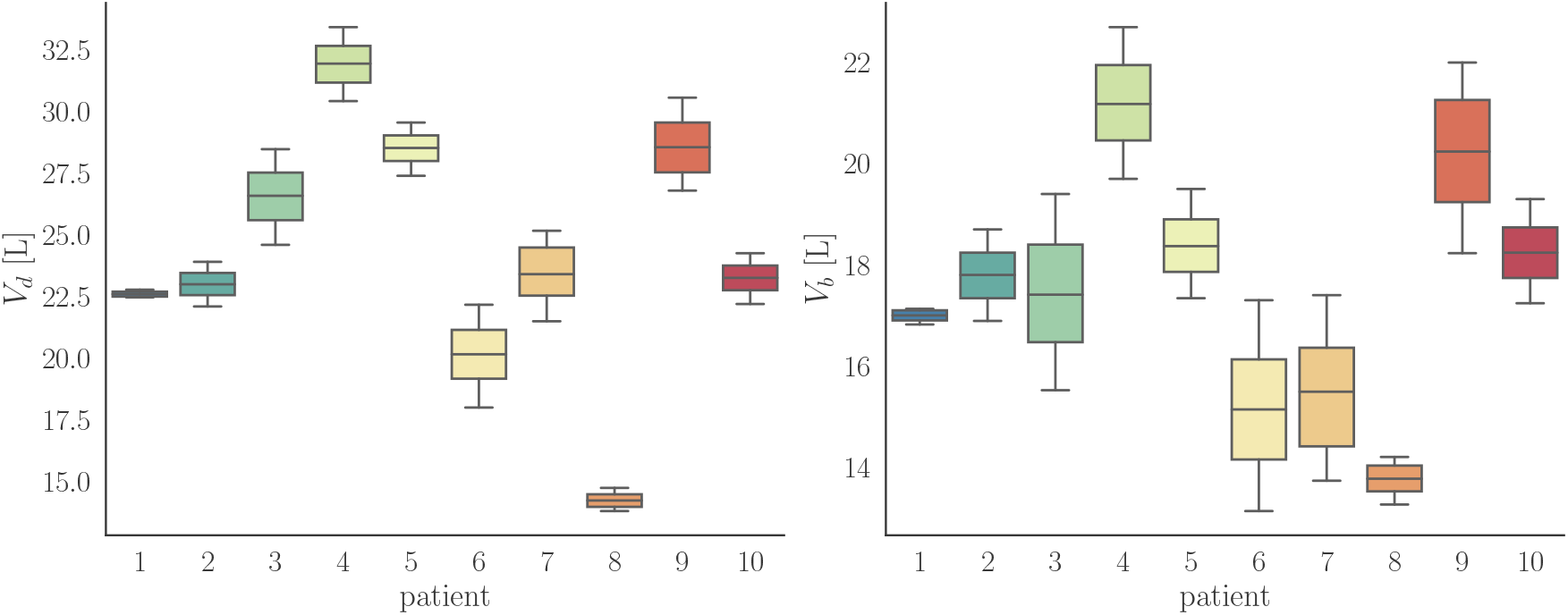
Boxplots of the measured parameters *V*_*b*_ and *V*_*d*_ for the MP sessions in Figure 3.

## B Tables

**Table B.1:**
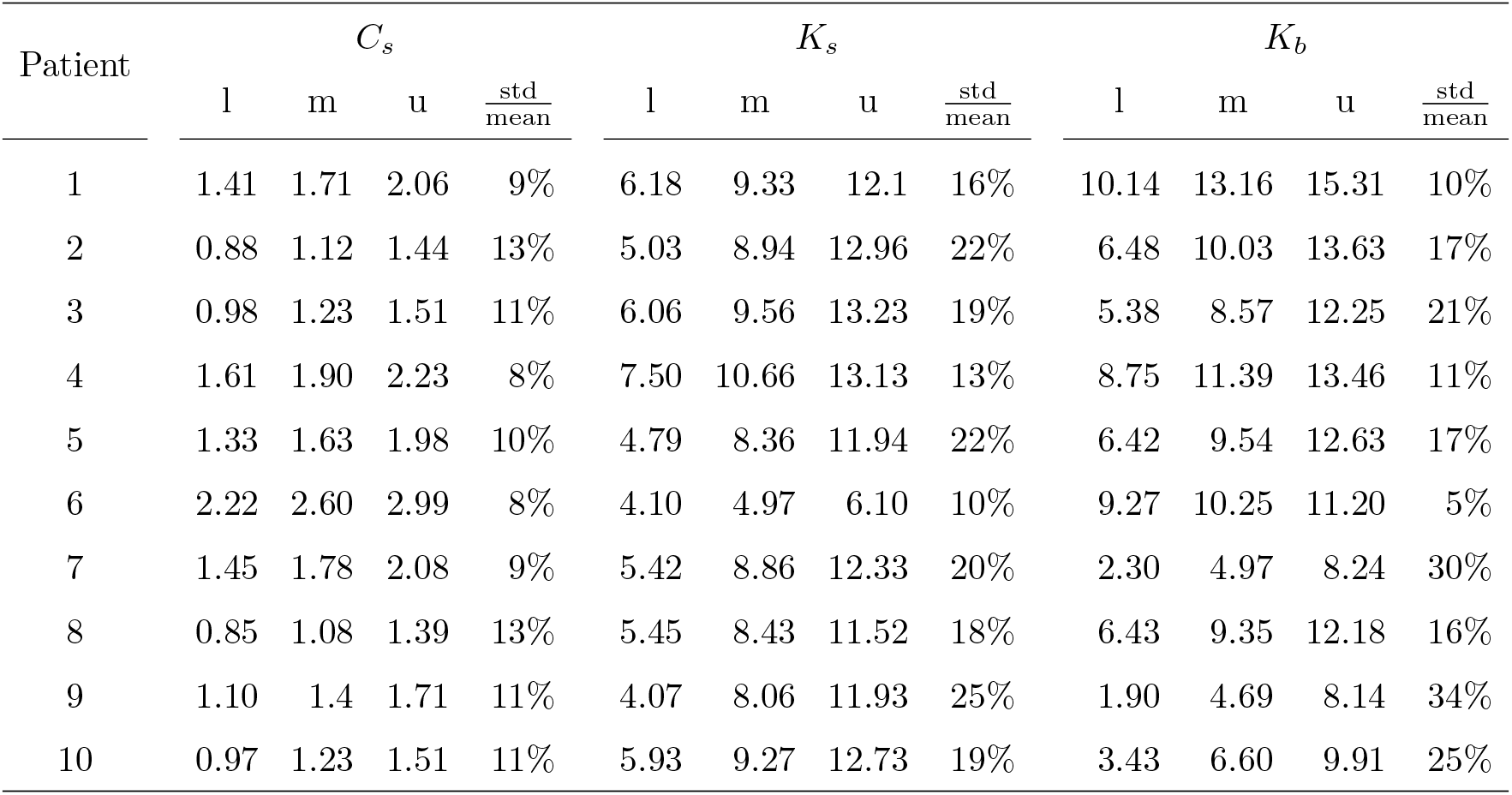
Median (m), lower (l) and upper (u) 95% CI and relative standard deviation 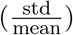 for the parameters for the SP estimation.

**Table B.2:** Median (m), lower (l) and upper (u) 95% CI and relative standard deviation 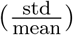 for the parameters for MP estimation.

| Patient | $C_s$ | | | | $K_s$ | | | | $K_b$ | | | |
| --- | --- | --- | --- | --- | --- | --- | --- | --- | --- | --- | --- | --- |
| | l | m | u | $\frac{\text{std}}{\text{mean}}$ | l | m | u | $\frac{\text{std}}{\text{mean}}$ | l | m | u | $\frac{\text{std}}{\text{mean}}$ |
| 1 | 1.41 | 1.57 | 2.13 | 12% | 2.30 | 6.98 | 11.97 | 38% | 5.82 | 7.08 | 8.73 | 10% |
| 2 | 1.05 | 1.15 | 1.37 | 7% | 4.38 | 7.80 | 11.58 | 24% | 7.39 | 8.47 | 9.78 | 7% |
| 3 | 1.26 | 1.40 | 1.62 | 7% | 4.96 | 7.78 | 11.22 | 21% | 6.48 | 7.78 | 9.41 | 10% |
| 4 | 1.29 | 1.45 | 1.71 | 7% | 4.50 | 7.46 | 11.06 | 23% | 7.14 | 8.49 | 10.16 | 9% |
| 5 | 1.31 | 1.42 | 1.65 | 6% | 4.45 | 7.23 | 10.21 | 22% | 9.43 | 10.07 | 10.79 | 3% |
| 6 | 1.23 | 1.37 | 1.65 | 8% | 3.71 | 7.13 | 11.06 | 26% | 6.04 | 7.50 | 9.43 | 11% |
| 7 | 1.50 | 1.64 | 1.91 | 6% | 4.22 | 7.89 | 11.86 | 24% | 5.84 | 7.12 | 8.75 | 11% |
| 8 | 0.90 | 0.99 | 1.19 | 7% | 3.07 | 7.36 | 11.45 | 28% | 3.22 | 4.34 | 6.9 | 20% |
| 9 | 1.16 | 1.28 | 1.44 | 6% | 5.95 | 9.03 | 12.63 | 19% | 4.96 | 5.98 | 7.43 | 10% |
| 10 | 1.2 | 1.32 | 1.54 | 7% | 4.40 | 7.11 | 10.66 | 23% | 6.54 | 7.16 | 7.88 | 5% |

**Table B.3:** Median (m), lower (l) and upper (u) 95% CI and relative standard deviation 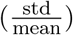 for the parameters for CP estimation.

| Patient | $C_s$ | | | | $K_s$ | | | | $K_b$ | | | |
| --- | --- | --- | --- | --- | --- | --- | --- | --- | --- | --- | --- | --- |
| | l | m | u | $\frac{\text{std}}{\text{mean}}$ | l | m | u | $\frac{\text{std}}{\text{mean}}$ | l | m | u | $\frac{\text{std}}{\text{mean}}$ |
| 1 | 1.37 | 1.51 | 1.91 | 8% | 3.19 | 7.26 | 11.34 | 28% | 7.07 | 8.27 | 9.78 | 8% |
| 2 | 1.03 | 1.11 | 1.27 | 5% | 5.44 | 8.45 | 11.72 | 19% | 8.2 | 9.30 | 10.51 | 6% |
| 3 | 1.24 | 1.38 | 1.60 | 7% | 5.11 | 7.70 | 11.14 | 19% | 7.23 | 8.38 | 9.74 | 8% |
| 4 | 1.34 | 1.51 | 1.79 | 8% | 4.30 | 7.25 | 11.14 | 24% | 6.57 | 7.59 | 8.79 | 7% |
| 5 | 1.26 | 1.34 | 1.50 | 5% | 6.72 | 10.48 | 13.93 | 18% | 8.20 | 9.09 | 10.05 | 5% |
| 6 | 1.37 | 1.66 | 2.24 | 14% | 2.02 | 4.21 | 8.41 | 39% | 5.67 | 6.70 | 7.86 | 8% |
| 7 | 1.55 | 1.67 | 1.86 | 5% | 5.89 | 9.20 | 12.74 | 19% | 4.82 | 5.81 | 7.08 | 10% |
| 8 | 0.86 | 0.96 | 1.13 | 7% | 3.39 | 7.09 | 11.05 | 27% | 4.27 | 5.64 | 7.59 | 16% |
| 9 | 1.22 | 1.33 | 1.52 | 6% | 5.42 | 8.54 | 12.28 | 20% | 4.59 | 5.36 | 6.27 | 9% |
| 10 | 1.17 | 1.24 | 1.36 | 4% | 6.70 | 9.62 | 12.62 | 16% | 6.28 | 6.97 | 7.68 | 5% |

**Table B.4:**
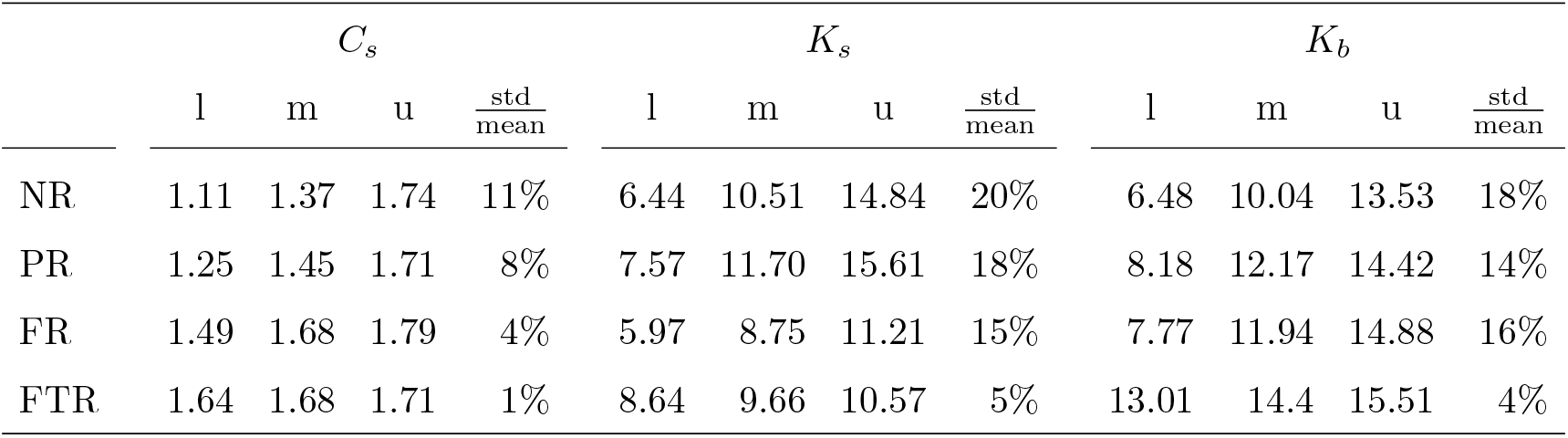
Median (m), lower (l) and upper (u) 95% CI and relative standard deviation 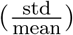 for the relapse data.

## C Correlation plots for SP and MP

**Figure C.1:**
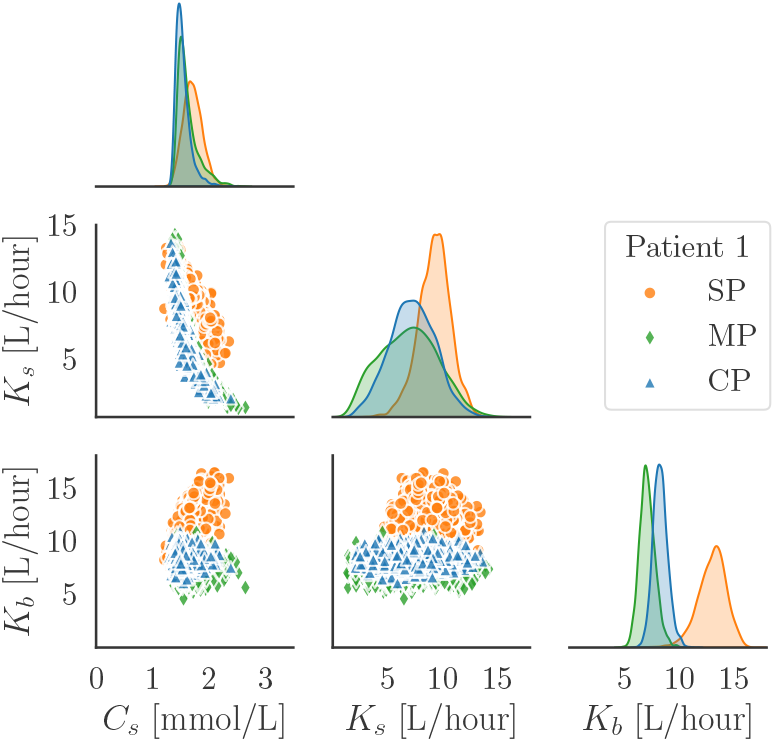
Correlation and posterior density for the patient 1.

**Figure C.2:**
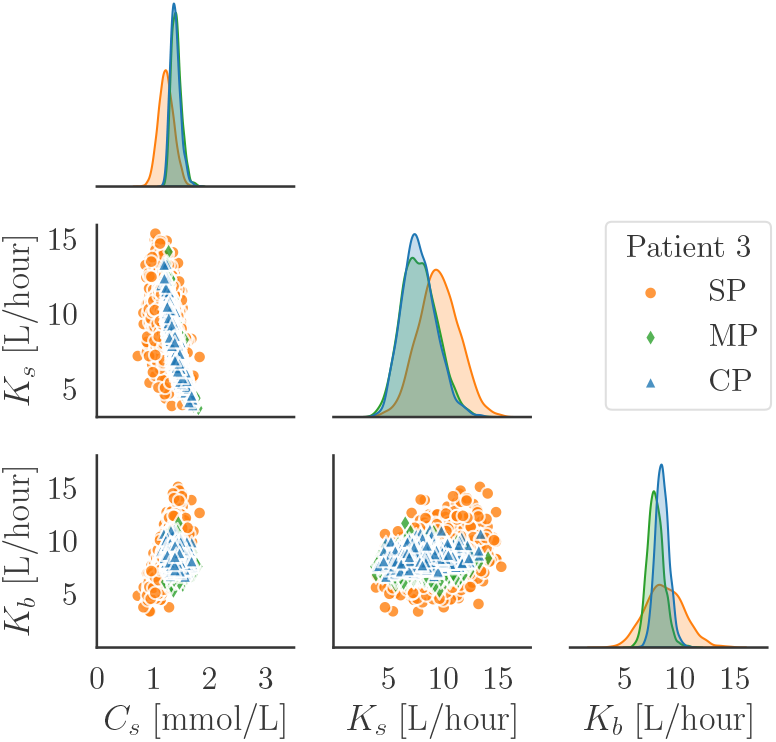
Correlation and posterior density for the patient 3.

**Figure C.3:**
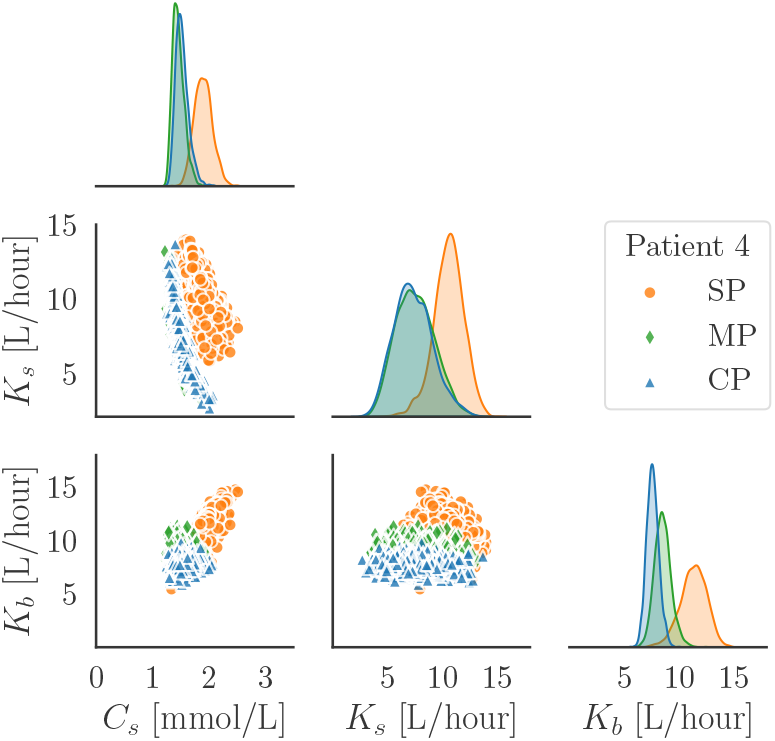
Correlation and posterior density for the patient 4.

**Figure C.4:**
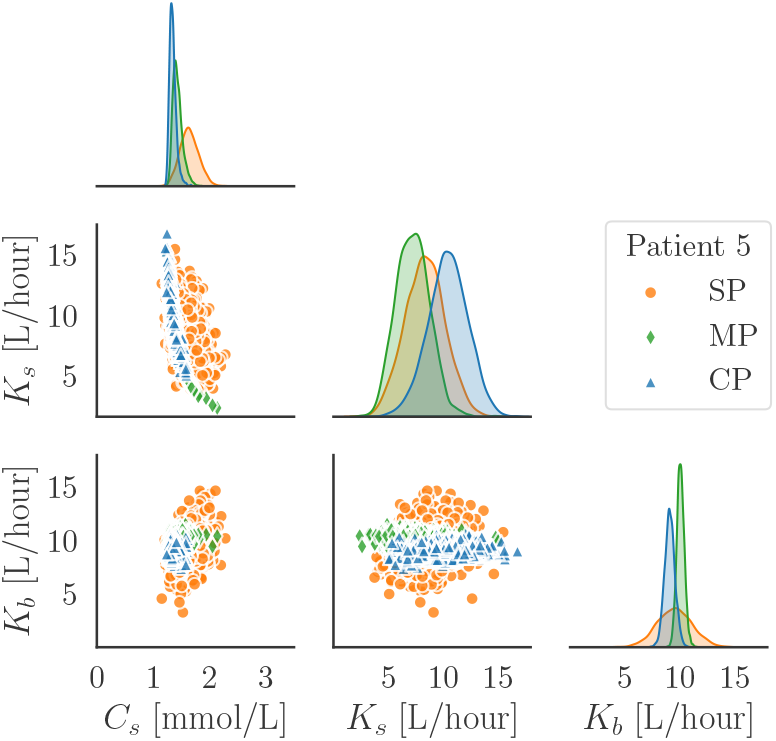
Correlation and posterior density for the patient 5.

**Figure C.5:**
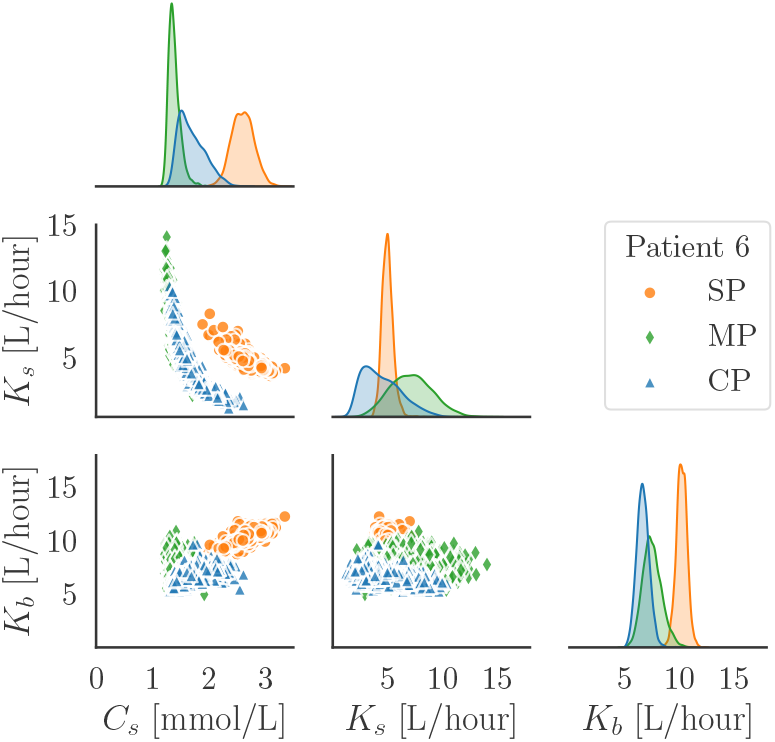
Correlation and posterior density for the patient 6.

**Figure C.6:**
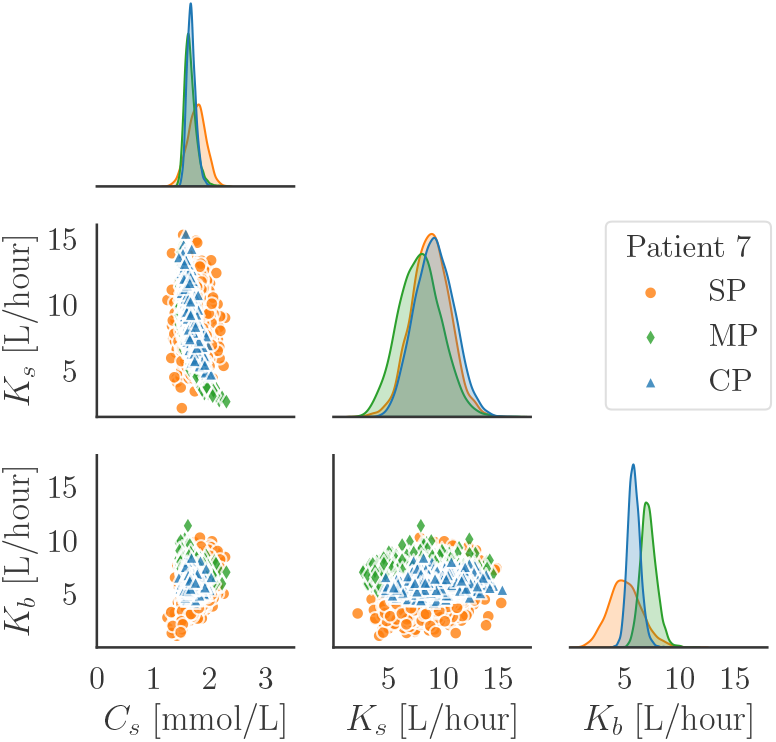
Correlation and posterior density for the patient 7.

**Figure C.7:**
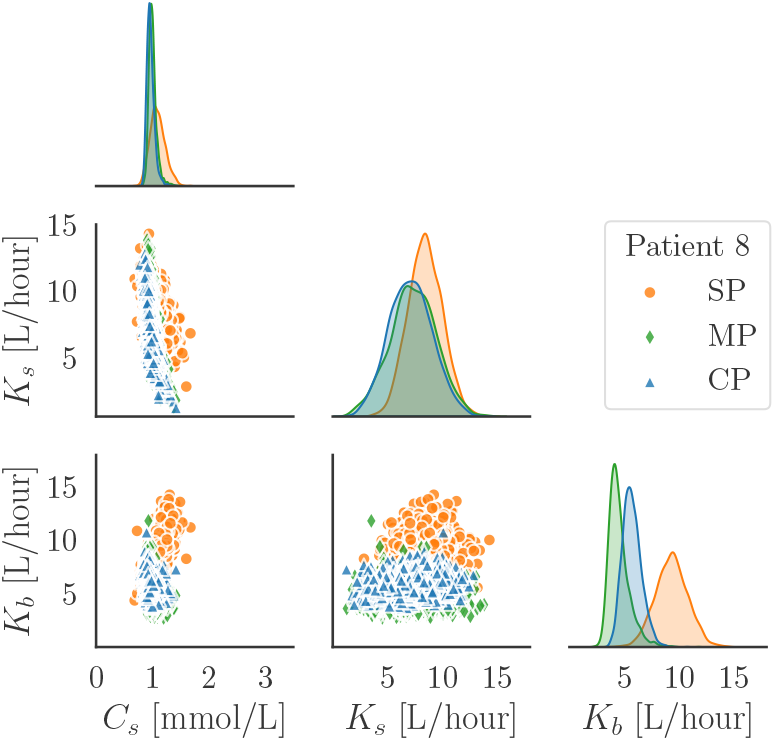
Correlation and posterior density for the patient 8.

**Figure C.8:**
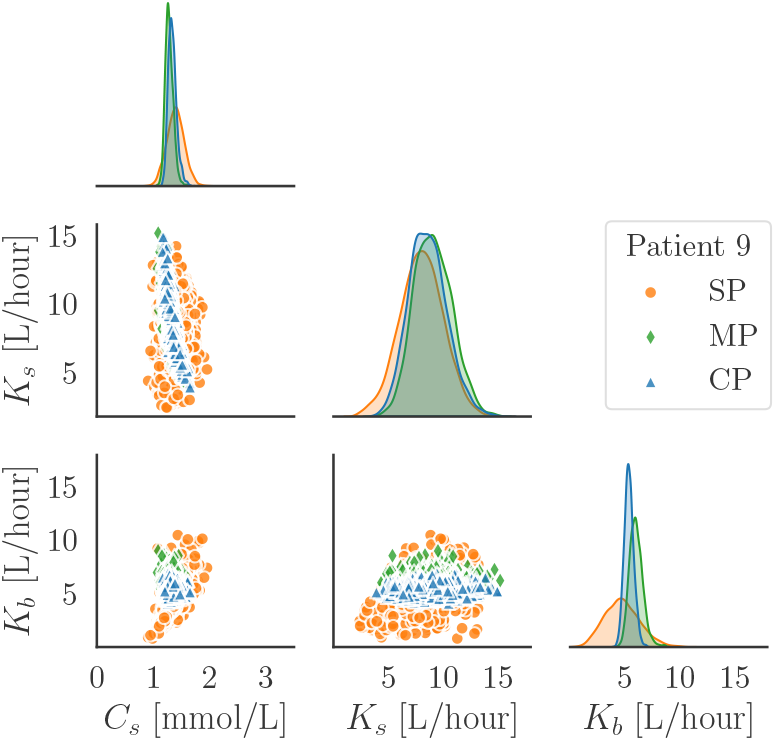
Correlation and posterior density for the patient 9.

**Figure C.9:**
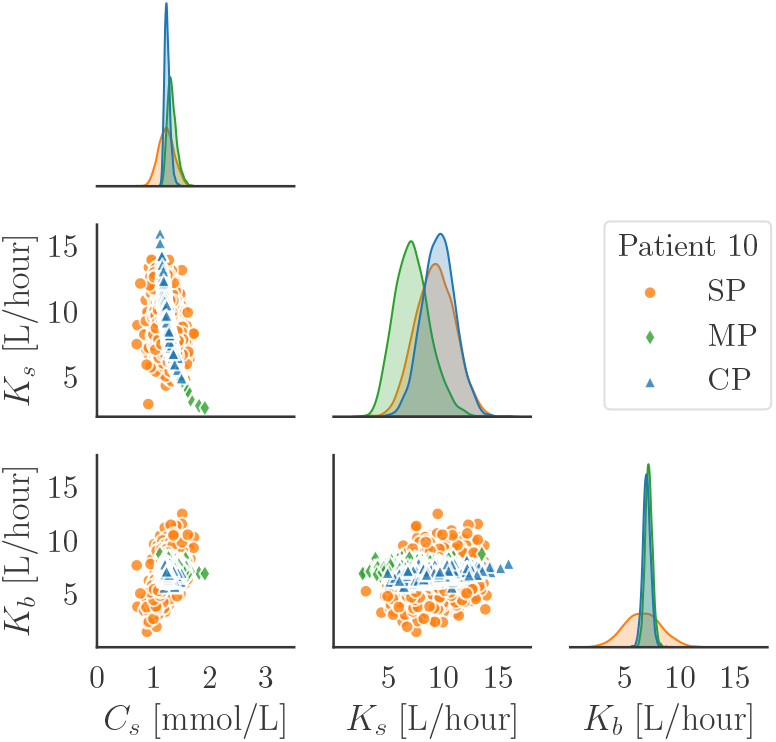
Correlation and posterior density for the patient 10.

